# Mapping nationally and globally at-risk species to identify hotspots for (and gaps in) conservation

**DOI:** 10.1101/2021.11.29.470436

**Authors:** Marie E Hardouin, Anna L Hargreaves

## Abstract

Protecting habitat of species-at-risk is critical to their recovery, but can be contentious. For example, protecting species that are locally imperilled but globally common (e.g. species that only occur in a jurisdiction at the edge of their geographic range) is often thought to distract from protecting globally-imperilled species. However, such perceived trade-offs are based on the assumption that threatened groups have little spatial overlap, which is rarely quantified. Here, we compile range maps of terrestrial species-at-risk in Canada to assess the geographic overlap of nationally and globally at-risk species with each other, among taxonomic groups, and with protected areas. While many nationally-at-risk taxa only occurred in Canada at their northern range edge (median=4% of range in Canada), nationally-at-risk species were not significantly more peripheral in Canada than globally-at-risk species. Further, 56% of hotspots of nationally-at-risk taxa were also hotspots of globally-at-risk taxa in Canada, undercutting the perceived trade-off in their protection. Hotspots of nationally-at-risk taxa also strongly overlapped with hotspots of individual taxonomic groups, though less so for mammals. While strong spatial overlap across threat levels and taxa should facilitate efficient habitat protection, <7% of the area in Canada’s at-risk hotspots is protected, and more than 70% of nationally and globally-at-risk species in Canada have <10% of their Canadian range protected. Our results counter the perception that protecting nationally vs. globally at-risk species are at odds, and identify critical areas to target as Canada strives to increase its protected areas and promote species-at-risk recovery.

## 1. INTRODUCTION

Habitat destruction is one of the primary threats to global biodiversity (Baillie et al., 2004). Indeed, the loss and degradation of habitat is the main threat to 85% of species listed in the International Union for the Conservation of Nature’s Red List of Threatened Species (IUCN, 2015) and a leading barrier to the recovery of imperilled species (Kerr & Deguise, 2004; Vellend, 2003). Protecting habitat through the establishment of protected areas is thus a crucial component of species-at-risk conservation. At the local scale, protected areas can alleviate land-conversion pressures on existing populations of species at risk, and at the landscape scale they can provide crucial habitat corridors that could facilitate species range shifts in response to climate change (Littlefield et al., 2019; Thomas et al., 2012).

While protected areas are a key conservation tool, it can be challenging to establish them in the areas most important for mitigating biodiversity loss. Habitat of species-at-risk often coincides with areas of high human population density or land use (Allan et al., 2019; Caissy et al., 2020), potentially generating mismatches between economic and conservation priorities. Combined with a lack of political will or funding (Watson et al., 2014), this can result in imperiled taxa being poorly represented within protected area networks (Venter et al., 2014). Given these challenges, optimizing which species-at-risk habitat is proposed for protection is crucial. A promising prioritization strategy is to identify ‘hotspots’ with particularly high taxonomic richness. This approach has been applied to specific taxonomic groups (Cameron & Hargreaves, 2020; Ceballos & Ehrlich, 2006; Fattorini et al., 2012; Orme et al., 2005), but it is unclear how well hotspots overlap among taxa. For instance, there is limited overlap in hotspots of vertebrates and plants in China (Xu et al., 2018) and of birds, mammals, and invertebrates in the UK (Prendergast et al., 1993). Poor hotspot overlap among taxa complicates prioritizing areas for conservation.

Even when hotspots align among taxa, prioritizing hotspots of at-risk species may be controversial when species are rare (and therefore deemed at-risk) in one jurisdiction but widespread elsewhere. Species can be rare in a jurisdiction simply because they only occur there at the edge of their geographic range (henceforth “peripheral” taxa; Bunnell et al., 2004). This can focus regional conservation efforts on range-edge populations of species that are globally secure, particularly in high-latitude countries that often contain the poleward range-edge of species more common toward the equator (Gibson et al., 2009). For instance, >75% of Canada’s nationally at-risk plants and Finland’s rare beetles are largely distributed outside these countries (Caissy et al., 2020; Komonen, 2007), and most US-state lists of threatened birds are dominated by taxa with low global risk (Wells et al., 2010). Protecting locally imperilled species has therefore been accused of being ‘parochial’, and coming at the expense of protecting more endemic or globally imperilled species (Hunter & Hutchinson, 1994; Rodrigues & Gaston, 2002). However, whether such a trade-off really exists if often unknown. Hotspots of locally at-risk species can co-occur with hotspots of overall diversity (Cameron & Hargreaves, 2020), and the extent to which hotspots of locally-at-risk and globally-at-risk species co-occur has rarely if ever been formally assessed.

An excellent case study for exploring how well hotspots of at-risk taxa co-occur, particularly between nationally and globally at-risk species, is Canada. As the world’s second largest country and the northernmost country in the Americas, habitat protection in Canada is critical for the species already there and the many predicted to shift northward under climate change. Because many nationally-at-risk taxa are peripheral in Canada (though this has only been quantified for plants and mammals), critics have argued that prioritizing them for conservation detracts from conserving globally-at-risk taxa (Kraus et al., 2021; Raymond et al., 2018). However, the geographic overlap between nationally and globally at-risk species has not been quantified. Further, while we know that Canada’s nationally at-risk taxa cluster in southern Canada (Coristine et al., 2018; Gibson et al., 2009), it is unclear whether this is driven by species-rich taxonomic groups (e.g. vascular plants, which cluster south; Caissy et al., 2020), and therefore how well overall hotspots of at-risk species would reflect hotspots of less species-rich groups (e.g. mammals, which have northwestern clusters; Cameron & Hargreaves, 2020). Identifying conservation hotspots in Canada is also particularly timely. Habitat loss threatens more than three-quarters of Canada’s nationally at-risk species (Woo-Durand et al., 2020), many of which have very little range overlap with protected areas (Caissy et al., 2020; Kraus & Hebb, 2020), and Canada has recently committed to protecting 25% of its land by 2025, up from the 12.5% conserved in 2020 (ECCC, 2021a; ECCC, 2021b).

Using range maps of Canada’s nationally and globally at-risk terrestrial taxa, we run three sets of analyses related to their conservation. *1) Hotspots*. We identify where species-at-risk are concentrated in Canada, and ask (Q1): How well do hotspots overlap among taxonomic groups, and between nationally and globally at-risk species? We predicted that most hotspots of nationally at-risk species would occur in southern Canada, where Canada’s biodiversity, human population, and human land use all cluster (Coristine et al., 2018; Kraus & Hebb, 2020). We did not have specific predictions about overlap given the distinct distributions of nationally at-ris plants and mammals (Caissy et al., 2020; Cameron & Hargreaves, 2020) and general sense that Canada’s globally-at-risk species are often in the far north. *2) Peripherality*. We quantify the proportion of each at-risk taxon’s range in Canada, and ask (Q2): Does peripherality (the extent to which taxa only occur in Canada at their range edge) vary among threat statuses and taxonomic groups, and between nationally and globally at-risk species? We expected that nationally-at-risk taxa would be mostly peripheral, and significantly more peripheral than globally-at-risk taxa, since national conservation assessment focuses on Canadian range area whereas the IUCN assesses global range area (COSEWIC, 2019; IUCN, 2012). *3) Habitat protection*. (Q3) How well is Canada protecting at-risk taxa? As many nationally at-risk species occur in highly human-modified southern Canada (Coristine et al., 2018; Coristine & Kerr, 2011), we were not optimistic about overlap with protected areas.

## 2. METHODS

Eligibility for protection under Canada’s Species at Risk Act (SARA) is determined by the Committee on the Status of Endangered Wildlife in Canada (COSEWIC). COSEWIC has seven subcommittees dedicated to assessing nine terrestrial taxonomic groups (Table 1) using quantitative criteria established by the IUCN. Taxa can be species, subspecies or populations. COSEWIC first prioritizes which taxa to assess, and at this stage prioritizes taxa likely to become globally extinct (SARA 15.1b). For prioritized taxa, COSEWIC commissions, reviews and approves an assessment report based on the ‘best biological information’ available (SARA, 2002), then uses the report to recommend a threat status. COSEWIC initially considers only Canadian populations, then reviews whether adjustments are warranted given the likelihood of demographic rescue from populations outside Canada (Environment and Climate Change Canada, 2017). Here we consider species assessed as nationally ‘at-risk’, i.e. Special Concern (may become threatened or endangered); Threatened (likely to become endangered if threats are not mitigated); or Endangered (facing imminent extirpation or extinction), and ignore taxa deemed Extirpated, Extinct, or Data deficient (i.e. no current Canadian range or reliable range map) or Not at risk. The final decision to list and protect taxa under SARA rests with the federal government after considering the socioeconomic implications (SARA, 2002). We use COSEWIC rather than SARA designations as they more closely reflect biology.

**Table 1.** Sample sizes for at-risk terrestrial taxa in Canada. Nationally at-risk taxa are those with a COSEWIC status of Special Concern, Threatened or Endangered. Globally at-risk taxa are those with an IUCN status of Vulnerable, Endangered or Critically Endangered. Range maps in C and D were obtained from COSEWIC for all taxa except birds, whose maps were obtained from BirdLife International and so are independent of sample sizes in A and B. C, D and G indicate the final sample sizes for Questions 1, 2, and 3. Missing Canadian range maps for globally at-risk species in F were obtained by digitizing COSEWIC maps when available.

|  | Nationally at-risk Taxa |  |  |  | Globally at-risk Taxa |  |  |
| --- | --- | --- | --- | --- | --- | --- | --- |
|  | A. Populations with a COSEWIC status | B. Taxa in A with a COSEWIC range map* | C. Taxa with a digitized Canadian range map (Q1, Q3**) | D. Taxa in C with a digitized global range map (Q2)*** | E. Taxa in Canada with an IUCN status | F. Taxa with an IUCN range polygon | G. Taxa with a digitized range map (Q2) |
| Birds | 76 | NA | 56 | 54 | 10 | 10 | 10 |
| Mammals | 42 | 41 | 37 | 36 | 7 | 7 | 7 |
| Reptiles | 41 | 34 | 32 | 26 | 3 | 1 | 3 |
| Amphibians | 27 | 24 | 21 | 17 | 0 | 0 | 0 |
| Arthropods | 70 | 66 | 60 | 54 | 12 | 6 | 9 |
| Molluscs | 40 | 39 | 34 | 32 | 17 | 14 | 15 |
| Vascular Plants | 207 | 200 | 184 | 171 | 21 | 15 | 17 |
| Mosses | 20 | 20 | 17 | 10 | 0 | 0 | 0 |
| Lichens | 23 | 23 | 21 | 14 | 6 | 2 | 3 |
| Total | 546 | NA | 462 | 414 | 76 | 55 | 64 |

\* If populations in A) were from the same species and only one range map was provided for that species, they were merged in B). \*\* Q3: analyses of hotspot overlap with protected areas uses $n$ in C; analyses of range overlap with protected areas by threat status has slightly lower $n$ , as populations of the same species with different threat statuses had to be excluded. \*\*\* If the populations merged in B) did not have the same COSEWIC status, they were excluded.

### 2.1 Data Collection

#### 2.1.1 COSEWIC Range Maps

COSEWIC assessment reports are publicly available as PDFs and generally include a range map of the assessed species (Government of Canada, 2021). For all taxa except birds (see 2.1.2) we updated a database developed by Pascale Caissy (see Caissy et al., 2020) by digitizing range maps in COSEWIC reports using the Quantum GIS 3.16 geographic information software (QGIS Development Team, 2021). We digitized the global range maps when possible, but 34 taxa had global range maps that were not properly digitizable (maps were incomplete, imprecise, or in a projection that could not be digitized), but Canadian range maps that were. For these taxa we digitized Canadian ranges only. These taxa were excluded from peripherality analyses (Q2) but included in the hotspot and protected area analyses (Q1, Q3) for which only the Canadian ranges were needed (Table 1).

To digitize COSEWIC maps, we first georeferenced maps by associating ≥15 (generally 40-60) specific points on the PDF maps with their corresponding coordinates on a base map of the world; these points were clear landmarks such as sharp land mass corners, waterbody edges, roads, or jurisdictional boundaries. Maps then underwent a thin plane spline transformation to project them in the World Geodetic System 1984 projection (WGS 84, EPSG:4326), a common global latitude and longitude-based coordinate reference system (Caissy et al., 2020; Kennedy & Kopp, 2000).

When ranges were already represented as polygons, we created a polygon shapefile by tracing the outline by hand; for ranges represented as point occurrences, we created a point shapefile by adding each point. If there were at least 4 points in the Americas, we generated a convex polygon (the smallest polygon shape encompassing all the points), which generally matched the taxon’s extent of occupancy as described in the text of COSEWIC assessment reports. If there were <4 point occurrences and we were unable to generate a convex polygon (12 taxa), we generated a buffer around each point instead. We set the buffer distance for each taxon to ensure that the taxon’s total buffered area would be roughly equivalent to the extent of occupancy reported in its COSEWIC report. We cut out the Great Lakes area from all range maps except for mollusc taxa that were present in these lakes.

#### 2.1.2 BirdLife Range Maps

Most bird species with a COSEWIC assessment are migratory, so their range maps had to be divided between their resident, breeding, non-breeding, and passage (e.g. migratory stopovers) ranges to meaningfully compare their Canadian and global distributions. We obtained range maps with this information from BirdLife International, who provided us with an ESRI File Geodatabase containing a separate distribution polygon shapefile for each seasonal range of each species (BirdLife International and Handbook of the Birds of the World, 2019). From these, we retained only ranges that overlapped a polygon shapefile of Canada (in QGIS3.16; QGIS Development Team, 2021). For species that spend multiple seasons in Canada (e.g. both breeding and passage) we merged the seasonal polygons and used this merged shapefile for analyses. Thus, we compared each species’ Canadian range with their corresponding seasonal global range. For instance, if a species only uses land in Canada during its breeding and passage season, we compared its Canadian breeding + passage range to its global breeding + passage range.

#### 2.1.3 IUCN Range Maps

To map globally at-risk species in Canada, we searched the IUCN Red List of Threatened Species database (IUCN, 2021) for species in the categories Vulnerable, Endangered or Critically Endangered (i.e. ‘at-risk’), whose ‘land region’ included Canada, and assessment scope = “global”. IUCN assessment reports are publicly available as PDFs, and most have corresponding polygon shapefiles of the species’ range. We downloaded these polygons, excluding 9 species whose IUCN reports were flagged as needing updating and 4 species whose Canadian ranges were defined as “vagrant” and did not appear in their global range maps. For each species we removed polygon areas defined by IUCN as “possibly extinct”. For species without a distribution polygon provided, we digitized the species’ range from its COSEWIC report if available (9 non-bird species; Table 1).

### 2.2 Data Extraction

Data extraction and analyses were done in R (version 4.1.0, R Core Team, 2021). To identify hotspots with unusually high concentrations of at-risk taxa, we projected each at-risk taxon’s digitized range map in the Albers equal area conic projection, which is commonly used for Canada (Kennedy & Kopp, 2000). We then overlaid these maps on a map of Canada divided into 100 × 100 km grid cells (1276 cells total) and counted the number of nationally and globally at-risk species in each grid cell. For each taxonomic grouping (the nine taxonomic groups of nationally-at-risk taxa, nationally-at-risk taxa overall, and globally-at-risk taxa overall) we identified cells with the highest species richness (“hotspots”), up to a maximum of 5% of the 1276 grid cells (max. 63 cells) (Cameron and Hargreaves 2020; adapted from Prendergast et al. (1993) Reid (1998)). E.g. cells containing 5 nationally-at-risk mammal taxa were not counted as mammal hotspots, as doing so would have resulted in >63 hotspot cells, leaving us with 55 mammal hotspot cells each with 6 or more at-risk mammal taxa. Thus the number of hotspot cells differed among taxonomic groups (42 to 62 hotspot cells per group).

For peripherality analyses, we projected digitized range maps in the World Mollweide equal area projection (WGS 84, ESRI:54009) to accommodate taxa with worldwide ranges. To assess peripherality, we quantified the proportion of each taxon’s western hemisphere range that occurs in Canada (i.e. the inverse of peripherality). We deemed western-hemisphere range to be a more biologically relevant denominator than global range, since the difficulty of moving between hemispheres makes it unlikely that populations on other continents would substantially impact Canadian populations. To calculate Canadian and western hemisphere range areas (km^2^), we intersected each digitized map with a Canada or western hemisphere (North, Central and South America) boundary map respectively (Natural Earth, 2020), thus cropping all maps to land only.

We calculated the proportion of each taxon’s Canadian range and the proportion of each hotspot that overlapped officially protected areas using the Canadian Protected and Conserved Areas Database (ECCC, 2021b). This publicly available database includes polygon shapefiles of “terrestrial protected areas” and “terrestrial areas with other effective area-based conservation measures” (ECCC, 2021b). We projected these protected area shapefiles in the Albers equal area conic projection. We then intersected the protected area shapefiles with 1) each at-risk taxon’s range polygon to calculate the km^2^ of protected habitat within each Canadian range; and 2) the map of Canada divided into 100 × 100 km grid cells from Q1 to calculate the km^2^ of protected habitat within each cell. We then divided the area of protected habitat within each grid cell by the grid cell’s total area, which was not always 100 × 100 km as some grid cells were cut by Canada’s borders.

### 2.3 Data Analyses

Analyses assessed differences among taxonomic groups for nationally-at-risk species, and compared nationally and globally at-risk species. We did not split analyses of globally-at-risk species by taxonomic group, due to small sample sizes in several groups (Table 1). Data used in analyses will be publicly archived on acceptance.

#### 2.3.1 Hotspot analyses

To assess hotspot overlap among pairs of groups (e.g. birds and mosses, nationally-at-risk and globally-at-risk), we counted the number of cells that were hotspots for both groups, then calculated the percentage of shared hotspots. As different taxonomic groups had different numbers of hotspots (see 2.2 above), we calculated overlap twice, one with each group as a denominator, then used their average. For example, birds (48 hotspots) and mosses (42 hotspots) shared 6 hotspot cells, resulting in a mean % overlap of 13% (i.e. the mean of 6/48 × 100 and 6/42 × 100).

#### 2.3.2 Peripherality analyses

We tested whether the proportion of a taxon’s western hemisphere range in Canada (proportional response) varied among taxonomic groups and threat statuses using generalized linear models (GLMs) with a beta-regression error structure (*glmmTMB* package version 1.1.2; Brooks et al., 2017). Beta-regressions deal with proportional data that does not result from a binomial process (such as percent areas), but cannot handle 0s and 1s. To deal with endemic species (i.e. range proportion in Canada = 1), we transformed all ‘range proportion in Canada’ values according to the equation (Douma & Weedon, 2019):

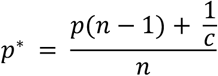

where *p*^*^ is the proportion (‘range proportion in Canada’), *n* is the total number of observations in the dataset (number of taxa) and *c* is the number of categories (2, i.e. range proportion in Canada and range proportion outside Canada).

First, we tested whether the ‘range proportion in Canada’ differed among COSEWIC statuses and/or taxonomic groups of nationally-at-risk taxa. Each nationally-at-risk taxon contributed one data point (sample sizes in Table 1D) so we used a generalized linear model (*Model 1*: prop range in Canada ∼ COSEWIC status x taxonomic group). We grouped vascular plants and mosses together (‘plants’) due to the small number of at-risk mosses (10 taxa; Table 1). Here and for all beta-regression models (see below) we tested the significance of fixed effects using likelihood ratio tests, which compare models with and without the fixed effect, compared to a χ^2^ distribution (*anova* function in R). As the 2-way interaction was not significant (χ^2^_df14_ = 16.8, *P* = 0.26), we dropped it from the final model to facilitate interpretation of main effects. For significant main effects, we used least-squared mean contrasts to determine which taxonomic groups or threat statuses differed from each other (*lsmeans* package version 2.30.0; Lenth, 2016).

Second, we tested the widely-held assumption that nationally-at-risk species are more peripheral in Canada than globally-at-risk species. We ran two models, each with one fixed effect, ‘threat status’. In *Model 2*, threat status had 6 categories (3 from COSEWIC and 3 from IUCN), to test whether peripherality varied with level of endangerment. In *Model 3*, we collapsed these into 2 categories ‘nationally-at-risk’ or ‘globally-at-risk’. In both models, each nationally-at-risk taxon and each globally-at-risk taxon contributed one data point (sample sizes in Table 1D&G). Because 42 taxa are considered at-risk by both COSEWIC and IUCN, and therefore occur in the data twice, we used a generalized linear mixed model with a random intercept for ‘species’ (prop range in Canada ∼ threat status + (1| species)). Significance of ‘threat status’ and differences among statuses were determined as in Model 1.

#### 2.3.3 Protected areas analyses

We tested how well hotspots of at-risk species were protected compared to less species-rich cells using beta regression GLMs. The response (proportion of a cell protected) was transformed as in 2.3.2 to deal with 0s and 1s (none vs. all of a cell protected). We tested whether the proportion of a grid cell that was protected (response) differed depending on whether or not the cell was a hotspot (binary predictor; model structure = prop cell protected ∼ hotspot status), with one model for nationally-at-risk hotspots (*Model 4*) and one for globally-at-risk hotspots (*Model 5*). Second, we tested whether the proportion of the cell that was protected varied with the number of at-risk taxa in the grid cell. Visual examination of the data suggested that this relationship might vary between hotspot and non-hotspot cells, so we included an interaction with hotspot status (prop cell protected ∼ # at-risk taxa x hotspot status). While this model also assessed the effect of hotspot status, we derive that effect from Models 4 and 5 as hotspot status is related to # at-risk taxa. We again ran one model for nationally-at-risk taxa (*Model 6)* and one for globally-at-risk taxa (*Model 7)*. We obtained backtransformed fit lines and 95% confidence intervals using the *visreg* package (version 2.7.0, Breheny & Burchett, 2017).

Finally, we tested how well ranges of at-risk species were being protected and whether this varied with threat status. We ran one beta regression model for nationally-at-risk taxa (*Model 8:* prop Canadian range protected ∼ COSEWIC status) and one for globally-at-risk taxa (*Model 9:* prop Canadian range protected ∼ IUCN status).

## 3. RESULTS

### 3.1 Hotspots of at-risk taxa

In Canada, hotspots of nationally-at-risk taxa generally clustered along the southern border (Figure 1, top nine maps). This was true for most taxonomic groups, with the notable exceptions of at-risk mammals, whose diversity hotspots followed the western mountains, and at-risk lichens, whose hotspots were mostly coastal (Figure 1). When all of Canada’s nationally at-risk taxa in our data were combined (bottom left map in Figure 1), their hotspots clustered along Canada’s southern borders from western Canada and the prairies to the Great Lakes and St-Lawrence river regions.

**Figure 1.**
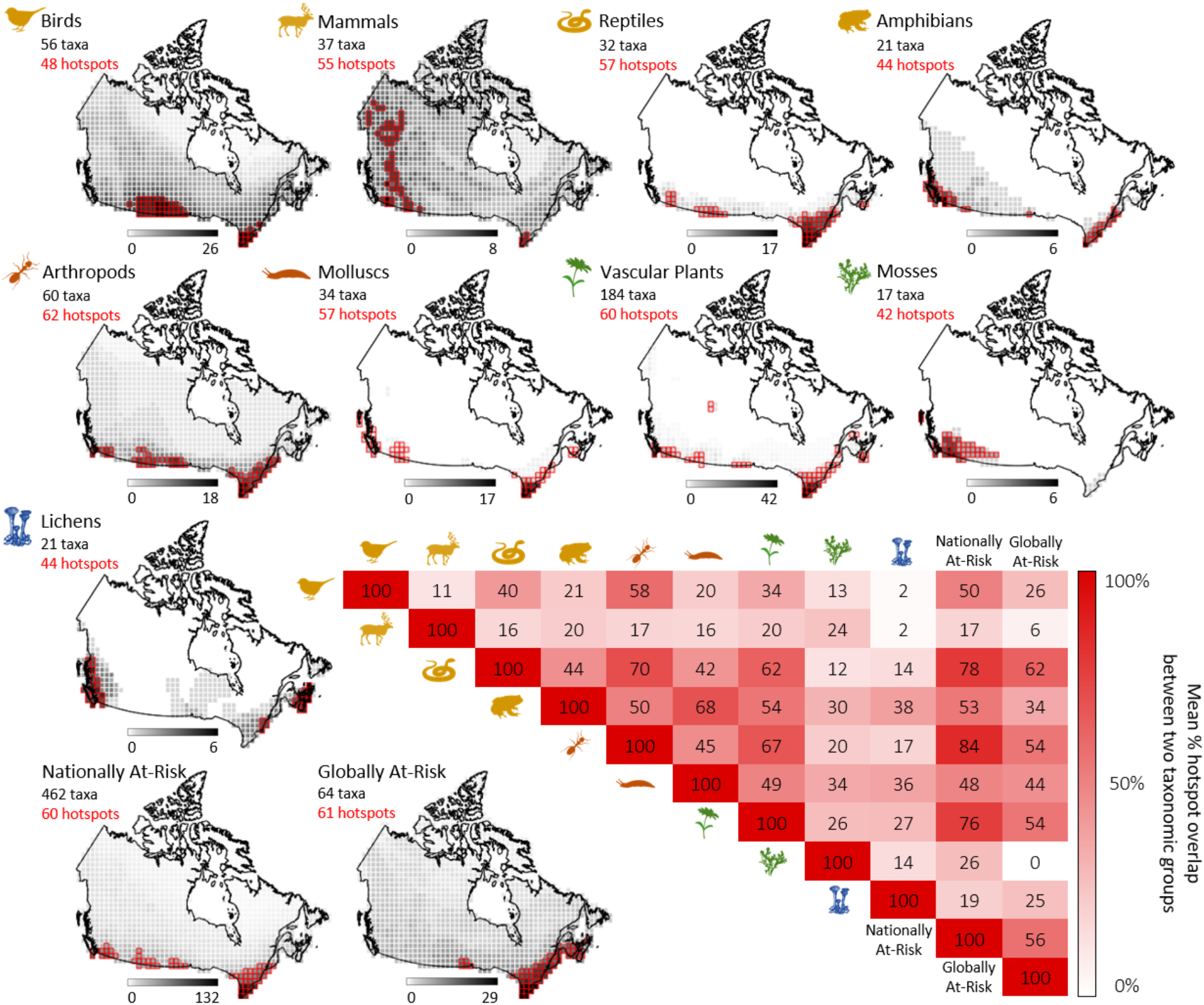
Hotspots of at-risk taxa in Canada. Maps are divided into 1276 100 × 100km grid cells, which are shaded in a greyscale according to the number of at-risk taxa they contain (greyscales differ among maps; legend below each map shows the maximum number of at-risk taxa per grid cell). Top three rows of maps show nationally-at-risk taxa separated by taxonomic group (yellow icons = vertebrates; orange = invertebrates; green = plants; blue = lichens). Bottom two maps show all nationally-at-risk (left) and globally-at-risk (right) taxa in Canada across taxonomic groups. Hotspot cells (i.e. those with the most at-risk taxa up to a maximum of 5% of the total grid cells) are outlined in red. The matrix of overlap (bottom right) indicates the percentage of hotspot cells that coincide for each pairwise combination.

Hotspots showed substantial taxonomic overlap. Taxonomic groups shared 31.5% of their hotspot cells on average (Figure 1). Overlap between hotspots of individual taxonomic groups and hotspots of nationally-at-risk taxa overall was even higher (mean = 50.1%). More than three quarters of the hotspots of at-risk reptiles, arthropods and vascular plants overlapped with nationally-at-risk hotspots (Figure 1). Hotspots of nationally-at-risk taxa contained up to 132 at-risk species, and 40% contained at-risk species from all 9 taxonomic groups. Together, the 4.4% of land in Canada included in nationally-at-risk hotspots was home to 80% of the 462 nationally-at-risk taxa in our data, including: 97% of the reptile taxa, 95% of amphibians, 92% of arthropods, 89% of birds, 85% of molluscs, 76% of plants, 65% of mosses, 65% of mammals, and 62% of lichens. While groups with the most at-risk species tended to have high overlap with national hotspots, overlap was not solely driven by richness. For example, there are twice as many at-risk vascular plant taxa in our data (184) than the next highest group (arthropods, 60 taxa), but arthropod hotspots had higher overlap with nationally-at-risk hotspots (84% vs. 70%; Figure 1).

Globally at-risk taxa in Canada showed a remarkably similar distribution to nationally-at-risk taxa (Figure 1, bottom two maps). Contrary to our expectation that globally-at-risk taxa might be distributed farther north, hotspots of globally-at-risk taxa also clustered along Canada’s southern border, but were almost entirely in eastern Canada from the Great Lakes to the southern Maritimes. More than half (56%) of globally at-risk hotspots were also hotspots of nationally at-risk taxa. Together, these 34 overlapping hotspots, representing only 2.4% of Canada’s land area, were home to 64% of the globally-at-risk taxa and 43% of the nationally-at-risk taxa in our data.

### 3.2 Peripherality of at-risk taxa in Canada

Most nationally-at-risk taxa only occur in Canada at the northern edge of their range (Figure 2A). While most taxonomic groups contained some endemic species (47 nationally-at-risk species had 100% of their western-hemisphere range in Canada, e.g. Vancouver Island Marmot, Banff Springs Snail, Magdalen Islands Grasshopper), 71% of nationally-at-risk taxa had less than 20% of their western-hemisphere range in Canada (mean = 23.8% of range in Canada, median = 4.4%). The extent to which taxa were peripheral in Canada varied among COSEWIC threat statuses (χ^2^_df2_ = 15.1, *P* < 0.001), and taxonomic groups (χ^2^_df7_ = 21.7, *P* = 0.003; *Model 1*). At-risk mammals had significantly more of their range in Canada (i.e. were less peripheral) than at-risk reptiles, amphibians, or plants (Figure 2C). The most imperilled taxa had significantly less of their range in Canada (i.e. were more peripheral) than the least imperilled at-risk taxa (Figure 2D).

**Figure 2.**
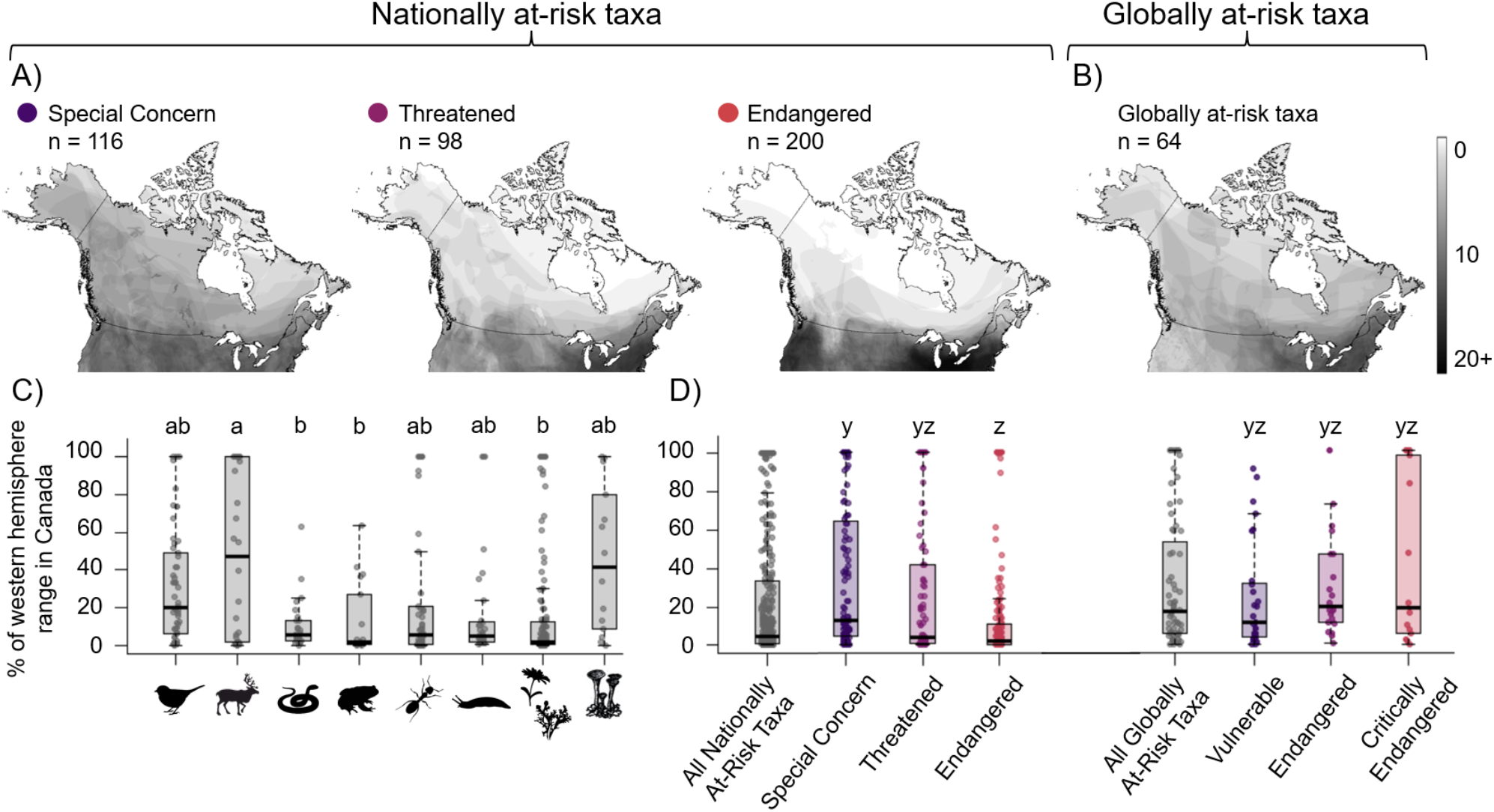
Geographic distribution and peripherality of at-risk taxa in Canada (that had digitizable Canadian and global range maps; see Table 1). Maps show Canadian ranges of **(A)** nationally-at-risk taxa, separated by their COSEWIC threat status, and **(B)** globally-at-risk taxa, combined across IUCN at-risk threat statuses. Coloured dots correspond to threat status, from least (purple) to most imperilled (red). **C)** Nationally-at-risk taxonomic groups differed in how peripheral they were; bars without shared letters are significantly different (*Model 1*). **D)** Nationally-at-risk taxa were not significantly more peripheral in Canada than globally-at-risk taxa when compared among the 6 threat statuses (*Model 2*; coloured bars) or between nationally and globally-at-risk taxa overall (*Model 3*; grey bars). Box plots show the median (thick middle line), 25^th^ and 75^th^ percentiles (box), 1.5x the interquartile range (whiskers), and raw data (horizontally jittered points).

Contrary to our predictions, many globally at-risk taxa also occurred in Canada only at their northern range edge. 54% of globally-at-risk taxa in Canada had less than 20% of their western hemisphere range in Canada (mean = 31.2% of range in Canada, median = 17.4%). All IUCN and COSEWIC threat categories contained both taxa with <5% of their range in Canada, and taxa that are endemic or almost endemic (IUCN Vulnerable taxa had up to 86.4% (for reindeer) of their range) in Canada. The extent to which taxa were peripheral in Canada varied significantly among the 6 threat statuses (*Model 2*: χ^2^_df5_ = 22.0, *P* = < 0.001; Figure 2D coloured bars), but only because nationally Endangered taxa had less of their range in Canada than Special Concern taxa, as found in *Model 1*. No category of nationally-at-risk taxa was more peripheral than any category of globally-at-risk taxa, nor did peripherality differ between nationally- and globally-at-risk taxa overall (*Model 3*; χ^2^_df1_= 2.8, *P* = 0.09; Figure 2D grey bars).

### 3.3 Habitat protection of at-risk taxa

Areas with particularly high concentrations of at-risk species (at-risk hotspots) are not particularly well protected. Whereas hotspots of nationally and globally at-risk taxa cluster in southern Canada, Canada’s largest expanses of protected habitat are further north (Figure 3). The percentage of a grid cell protected did not differ between cells that were hotspots of nationally-at-risk taxa (mean = 8.0 ± 12.3% of cell protected, median = 3.5%) and cells that were not (mean = 12.9 ± 22.6%, median = 2.4%; *Model 4*: χ^2^_df1_ = 1.2, *P* = 0.27), nor between cells that were hotspots of globally-at-risk taxa (mean = 4.7 ± 5.9%, median of 2.7%) and cells that were not (mean = 13.1 ± 22.6%, median = 2.5%; *Model 5*: χ^2^_df1_ = .51, *P* = 0.48). For both nationally and globally at-risk taxa, the percentage of a grid cell protected increased as the number of at-risk taxa in the cell increased, but only for non-hotspot cells (significant interaction # at-risk taxa x hotspot status: *Model 6*: χ^2^_df1_ = 32.7, *P* < 0.001; *Model 7:* χ^2^_df1_ = 18.9, *P* < 0.001; Figure 3C&D). Of the 1276 grid cells across Canada, 34% contained no protected areas, while 0.5% were fully protected. Only 6.8% of the land in nationally-at-risk hotspots was protected. Protection is even lower for land in hotspots of globally-at-risk taxa (5.1% protected), and lower still for areas that were hotspots for both nationally and globally-at-risk taxa (3.8% protected).

**Figure 3.**
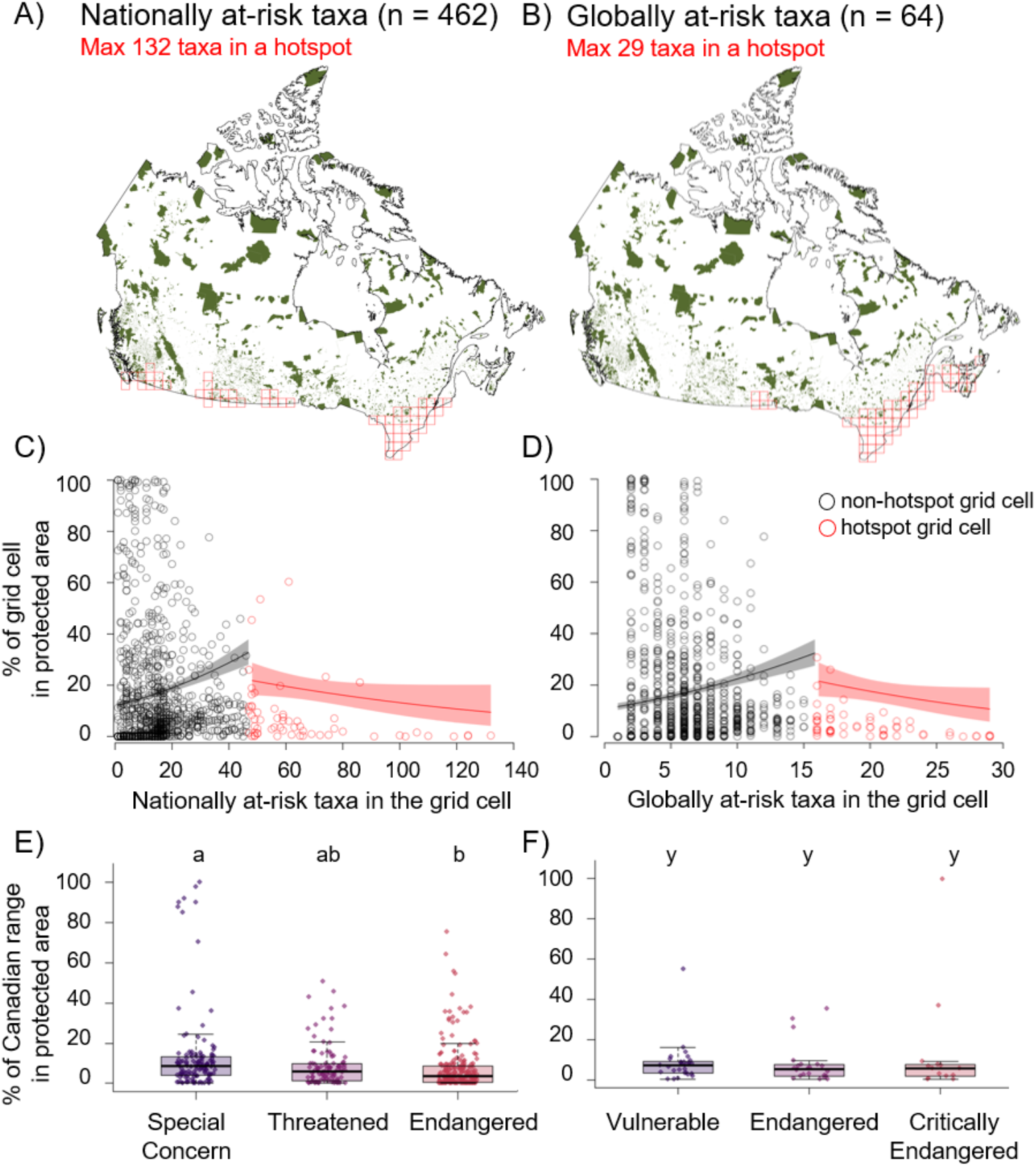
Habitat protection of nationally-at-risk taxa (left) and globally-at-risk taxa (right) in Canada. **A-B)** location of Canada’s publicly protected and conserved areas (green), and hotspots of at-risk taxa (from Figure 1) outlined in red. **C-D)** protection increased significantly with the number of at-risk taxa in a grid cell, but only for cells that were not hotspots of at-risk taxa (black). Each point represents one of the 1276 grid cells in Canada (raw data), trend lines and 95% confidence intervals are extracted from *Model 6* (C) and *Model 7* (D). **E-F)** Percentage of at-risk taxa’s Canadian range included in protected areas. Differing letters represent significant differences among COSEWIC threat statuses (*Model 8*; E) or IUCN threat statuses (*Model 9;* F). Sample sizes are in Table 1. Box plot formats are as in Figure 2.

The Canadian ranges of individual at-risk taxa were also poorly protected on average. Nationally and globally at-risk taxa in Canada had only 5.5% and 5.7% (median) of their Canadian range protected, respectively. Most nationally-at-risk taxa (74%) and globally at-risk taxa (83%) had less than 10% of their Canadian range protected. This is despite the fact that many at-risk taxa have small range areas in Canada (median range area in Canada = 12,157 km^2^ for nationally-at-risk taxa and 105,634 km^2^ for globally-at-risk taxa). Habitat protection differed among COSEWIC statuses, with the most imperilled taxa having *less* of their Canadian range protected than the least imperilled at-risk taxa (*Model 8*: χ^2^_df2_ = 15.3, *P* < 0.001; Figure 3E). Habitat protection was equally low among IUCN statuses (*Model 9*: χ^2^_df2_ = 0.39, *P* = 0.82; Figure 3F).

## 4. DISCUSSION

Our analyses of Canada’s at-risk terrestrial taxa support some preconceptions about their spatial distribution, but counter others. First, our results confirm that most species considered at-risk in Canada only occur in Canada at the northern edge of their range (Figure 2). This aligns with previous qualitative assessments of species-at-risk in Canada (Gibson et al. 2009), but had not been quantified across taxa before. However, our results do not support the perception that nationally-at-risk taxa are more peripheral than globally-at-risk taxa in Canada. While globally-at-risk species did have slightly more of their range in Canada on average, they also clustered toward the southern border and 54% had less than 20% of their range in Canada (Figure 2). These results underscore that conservation of at-risk species and biodiversity in Canada, as in many high latitude countries, is fundamentally linked to the conservation of range-edge populations.

Our results also challenge the perception that protecting nationally-at-risk species inherently involves a trade-off with protecting globally-at-risk species. As so many of Canada’s nationally-at-risk taxa clusters in the south due to populations at the edge of their range (Kraus et al., 2020), there has been significant debate about how to prioritize their conservation (Fraser 2000), and sustained criticisms that doing so comes at the cost of protecting globally-imperilled taxa (Jones & Fredricksen, 1999; Raymond et al., 2018). Our results suggest that at least in terms of habitat protection, this is a false dichotomy. More than half (56%) of the areas with the highest concentration of nationally-at-risk taxa in Canada were also hotspots of globally-at-risk species in Canada (Figure 1). Hotspots of nationally-at-risk taxa encompassed 64% of the globally at-risk taxa in our data, showing that protecting one threat group can significantly benefit the other.

Not only do hotspots of nationally-at-risk taxa contain many globally-at-risk taxa, they are also taxonomically diverse. A single nationally-at-risk hotspot contained up to 132 at-risk species (Figure 1), and all nationally-at-risk hotspots contained at-risk species from at least 6 taxonomic groups. However, nationally-at-risk hotspots did not represent at-risk mammals as well as they did other taxa. This is important as non-human mammals often garner disproportionate conservation interest from human mammals (Davies et al., 2019; Donaldson et al., 2017; Titley et al., 2017). Hotspots of nationally-at-risk mammals do contain disproportionately high overall mammal diversity (Cameron & Hargreaves, 2020), so conservation efforts in these areas are certainly worthwhile. However, our results caution that using at-risk mammals as charismatic flagship species will not necessarily focus conservation efforts toward the habitat most needed by other at-risk taxa. Protecting habitat in nationally-at-risk hotspots, on the other hand, can have a significant conservation benefit for at-risk mammals. Hotspots of nationally-at-risk species together contained >50% of the at-risk taxa in each taxonomic group, which is highly promising for planning diverse protected areas.

Strong overlap in hotspots of different taxonomic groups and threat levels should facilitate habitat protection to promote species-at-risk recovery, but we see little evidence that protected areas have so far been planned with species-at-risk in mind. The amount of land protected did increase with the number of at-risk species in non-hotspots cells (Figure 3), which is an improvement over a complete lack of relationship between protected areas and density of species at-risk 15 years ago (Deguise & Kerr 2006). However, this effect was slight, and most nationally and globally at-risk species have only a tiny fraction of their Canadian range protected (Figure 3). Areas that are hotspots for both nationally-at-risk and globally-at-risk taxa should be of the highest conservation priority, but only 3.8% of these areas are publicly protected. This shows a sharp disconnect between species at-risk policy and protected area designations.

Why is Canada not doing better at protecting species-at-risk within its borders? The disconnect could reflect delays and biases in identifying critical habitat or implementing recovery plans (Bird & Hodges, 2017; Findlay et al., 2009; Mooers et al., 2010), or lack of co-ordination between the disparate government departments responsible for species-at-risk recovery planning and protected area designations. It strongly suggests that protected area selection has so far been driven by objectives other than species-at-risk protection. While protected area objectives can include historical, scenic, or cultural significance (Benidickson, 2013), designation may also be influenced by minimizing resource exploitation conflicts, rather than optimizing species-at-risk conservation (Bolliger et al., 2020; Venter et al., 2017) as is the case in many protected area networks worldwide (Clancy et al., 2020; Venter et al., 2014). The difficulty of protecting habitat for species-at-risk species underscores the importance of identifying priority areas that can promote as many conservation objectives as possible. Our findings of high spatial overlap among at-risk taxa is great news: it means Canada’s recent commitment to protect more natural area could easily be used to promote species-at-risk recovery. Further, protecting natural landscapes in densely populated areas like southern Canada can also benefit the many people living there (Jiricka-Pürrer et al., 2019; White et al., 2019), who are increasingly seeking natural areas and greenspace (Bain, 2021; Parks Canada, 2019).

Of course, habitat protection generally happens at a smaller scale that the 100 × 100 km grid cells used in our hotspot analysis. This would be especially true for new protected areas developed in southern Canada where private land ownership is high. Since species rarely occupy an entire hotspot grid cell or even all the area within their distribution polygons (Rotenberry & Balasubramaniam, 2020), protecting a smaller subsection of habitat might not encompass a hotspot’s full suite of overlapping taxa. Even so, coarsely mapping Canada’s nationally and globally at-risk hotspots provides a useful starting point for future, more local-scale analyses to pinpoint priority areas. Further, patchy species distributions mean that small parks and protected areas can have a disproportionate benefit for individual species-at-risk.

Another reason to protect habitat in hotspots of species at risk, even if not all species in the region occur in the local protected area, is that species range shifts driven by climate change are already changing taxonomic overlap (Yong et al., 2016). Protecting land within the taxonomically diverse, at-risk hotspots we identified could offer habitat refuges to surrounding species whose ranges do not yet overlap that area. Northward range shifts are especially important given our finding that 71% and 54% of the nationally and globally at-risk taxa in Canada occur at their northern range edge. Protecting habitat in at-risk hotspots would not only protect a taxonomically diverse set of at-risk populations, but could also provide essential corridors for edge populations to shift toward higher latitudes and track a changing climate (Chen et al., 2011; Gibson et al., 2009). Protecting habitat within Canada’s at-risk hotspots is therefore crucial for Canada’s current and future biodiversity.

## 5. ACKNOWLEDGMENTS

We thank Pascale Caissy for spearheading and developing methods for digitizing COSEWIC range maps, and the Liber Ero Chair in Biodiversity Conservation at McGill University for financial support in developing this database. We also thank Pascale and Victor Cameron for helping MEH with further map digitization and quantifying spatial distribution hotspots, respectively, and the McGill Geographic Information Centre for help with geographic data. Funding for this project was provided by the Natural Sciences and Engineering Research Council of Canada (NSERC) through an Undergraduate Science Research Award to MEH and Discovery Grant to ALH.

## Notes

### Competing Interest Statement

The authors have declared no competing interest.

### Summary of Updates

doi to one of the reference papers has been corrected

## REFERENCES

Allan, J. R., Watson, J. E. M., Di Marco, M., O’Bryan, C. J., Possingham, H. P., Atkinson, S. C., & Venter, O. (2019). Hotspots of human impact on threatened terrestrial vertebrates. PLoS Biology, 17(3), e3000158. https://doi.org/10.1371/journal.pbio.3000158

Baillie, J. E. M., Hilton-Taylor, C., & Stuart, S. N. (2004). 2004 IUCN Red List of Threatened Species. A global species assessment. IUCN, Gland, Switzerland and Cambridge, UK. Xxiv + 191 pp

Bain, J. (2021). Visitor stats reveal how Canada’s parks and historic sites fared in 2020. National Parks Traveler. Available from https://www.nationalparkstraveler.org/2021/03/visitor-stats-reveal-how-canadas-parks-and-historic-sites-fared-2020

Benidickson, J. (2013). Legal framework for protected areas: Canada. IUCN-EPLP, 81, 1–35. https://doi.org/10.2139/ssrn.2296254

Bird, S. C., & Hodges, K. E. (2017). Environmental science & policy critical habitat designation for Canadian listed species : slow, biased, and incomplete. Environmental Science and Policy, 71, 1–8. https://doi.org/10.1016/j.envsci.2017.01.007

BirdLife International and Handbook of the Birds of the World. (2019). Bird species distribution maps of the world. Version 2019.1. Available from http://datazone.birdlife.org/species/requestdis

Bolliger, C. S., Raymond, C. V., Schuster, R., & Bennett, J. R. (2020). Spatial coverage of protection for terrestrial species under the Canadian Species at Risk Act. Ecoscience, 27(2), 141–147. https://doi.org/10.1080/11956860.2020.1741497

Breheny, P., & Burchett, W. (2017). Visualization of regression models using visreg. The R Journal, 9(2), 56–71.

Brooks, M. E., Kristensen, K., Benthem, K.J. Van, Magnusson, A., Berg, C. W., Nielsen, A., Skaug, H. J., Mächler, M., & Bolker, B. M. (2017). glmmTMB balances speed and flexibility among packages for zero-inflated generalized linear mixed modeling. The R Journal, 9(2), 378–400.

Bunnell, F. L., Campbell, R. W., & Squires, K. A. (2004). Conservation priorities for peripheral species : the example of British Columbia. Canadian Journal of Forestry Research, 34, 2240–2247. https://doi.org/10.1139/X04-102

Caissy, P., Klemet-N’Guessan, S., Jackiw, R., Eckert, C. G., & Hargreaves, A. L. (2020). High conservation priority of range-edge plant populations not matched by habitat protection or research effort. Biological Conservation, 249, 108732. https://doi.org/10.1016/j.biocon.2020.108732

Cameron, V., & Hargreaves, A. (2020). Spatial distribution and hotspots of mammals in Canada. Facets, 5, 1–12. https://doi.org/10.1139/facets-2020-0018

Ceballos, G., & Ehrlich, P. R. (2006). Global mammal distributions, biodiversity hotspots, and conservation. Proceedings of the National Academy of Sciences of the United States of America, 103(51), 19374–19379. https://doi.org/10.1073/pnas.0609334103

Chen, I. C., Hill, J. K., Ohlemüller, R., Roy, D. B., & Thomas, C. D. (2011). Rapid range shifts of species associated with high levels of climate warming. Science, 333(6045), 1024–1026. https://doi.org/10.1126/science.1206432

Clancy, N. G., Draper, J. P., Wolf, J. M., Abdulwahab, U. A., Pendleton, M. C., Brothers, S., Brahney, J., Weathered, J., & Hammill, E. (2020). Protecting endangered species in the USA requires both public and private land conservation. Scientific Reports, 1–8. https://doi.org/10.1038/s41598-020-68780-y

Coristine, L. E., Jacob, A. L., Schuster, R., Otto, S. P., Baron, N. E., Bennett, N. J., Bittick, S. J., Dey, C., Favaro, B., Ford, A., Nowlan, L., Orihel, D., Palen, W. J., Polfus, J. L., Shiffman, D. S., Venter, O., & Woodley, S. (2018). Informing Canada’s commitment to biodiversity conservation: a science-based framework to help guide protected areas designation through Target 1 and beyond. Facets, 3(1), 531–562. https://doi.org/10.1139/facets-2017-0102

Coristine, L. E., & Kerr, J. T. (2011). Habitat loss, climate change, and emerging conservation challenges in Canada 1. Canadian Journal of Zoology, 89(5), 435–451. https://doi.org/10.1139/Z11-023

COSEWIC. (2019). COSEWIC Assessment Process, Categories, and Guidelines.

Davies, T., Cowley, A., Bennie, J., Leyshon, C., Inger, R., Carter, H., Robinson, B., Duffy, J. P., Casalegno, S., Lambert, G., & Gaston, K. (2019). Correction: Popular interest in vertebrates does not reflect extinction risk and is associated with bias in conservation investment (PLoS ONE (2018) 13: 9 (e0203694) DOI: 10.1371/journal.pone.0203694). PLoS ONE, 14(2), 1– https://doi.org/10.1371/journal.pone.0212101

Deguise, I. E., & Kerr, J. T. (2006). Protected Areas and Prospects for Endangered Species Conservation in Canada. Conservation Biology, 20(1), 48–55. https://doi.org/10.1111/j.1523-1739.2005.00274.x

Donaldson, M. R., Burnett, N. J., Braun, D. C., Suski, C. D., Hinch, S. G., Cooke, S. J., & Kerr, J. T. (2017). Taxonomic bias and international biodiversity conservation research. Facets, 1(1), 105–113. https://doi.org/10.1139/facets-2016-0011

Douma, J. C., & Weedon, J. T. (2019). Analysing continuous proportions in ecology and evolution: a practical introduction to beta and Dirichlet regression. Methods in Ecology and Evolution, 10(9), 1412–1430. https://doi.org/10.1111/2041-210X.13234

Environment and Climate Change Canada. (2021a). Canada Target 1 challenge. Available from https://www.canada.ca/en/environment-climate-change/services/nature-legacy/canada-target-one-challenge.html

Environment and Climate Change Canada. (2021b). Canadian protected and conserved areas database. Available from https://www.canada.ca/en/environment-climate-change/services/national-wildlife-areas/protected-conserved-areas-database.html

Fattorini, S., Dennis, R. L. H., & Cook, L. M. (2012). Use of Cross-Taxon congruence for hotspot identification at a regional scale. PLoS ONE, 7(6), 1–6. https://doi.org/10.1371/journal.pone.0040018

Findlay, C. S., Elgie, S., Giles, B., & Burr, L. (2009). Species listing under Canada’s Species at Risk Act. Conservation Biology, 23(6), 1609–1617. https://doi.org/10.1111/j.1523-1739.2009.01255.x

Gibson, S. Y., Van Der Marel, R. C., & Starzomski, B. M. (2009). Climate change and conservation of leading-edge peripheral populations. Conservation Biology, 23(6), 1369–1373. https://doi.org/10.1111/j.1523-1739.2009.01375.x

Government of Canada. (2021). Species at risk public registry. Available from https://species-registry.canada.ca/index-en.html

Hunter, M. L., & Hutchinson, A. (1994). The virtues and shortcomings of parochialism: conserving species that are locally rare, but globally common. Conservation Biology, 8(4), 1163–1165. https://doi.org/10.1046/j.1523-1739.1994.08041163.x

IUCN. (2012). IUCN Red List categories and criteria: Version 3.1. Second edition. Gland, Switzerland and Cambridge, UK: IUCN. iv + 32pp.

IUCN. (2015). Conservation successes overshadowed by more species declines – IUCN Red List update | IUCN. Available from https://www.iucn.org/content/conservation-successes-overshadowed-more-species-declines---iucn-red-list-update

IUCN. (2021). The IUCN Red List of Threatened Species. Version 2021-1. Available from https://www.iucnredlist.org. Downloaded on 2021-05-08.

Jiricka-Pürrer, A., Tadini, V., Salak, B., Taczanowska, K., Tucki, A., & Senes, G. (2019). Do protected areas contribute to health and well-being? A cross-cultural comparison. International Journal of Environmental Research and Public Health, 16(7). https://doi.org/10.3390/ijerph16071172

Jones, L., & Fredricksen, L. (1999). Crying wolf? Public policy on endangered species in Canada. Critical Issues Bulletin, The Fraser Institute, 1–23.

Kearney, S. G., Adams, V. M., Fuller, R. A., Possingham, H. P., & Watson, J. E. M. (2020). Estimating the benefit of well-managed protected areas for threatened species conservation. Oryx, 54(2), 276–284. https://doi.org/10.1017/S0030605317001739

Kennedy, M., & Kopp, S. (2000). Understanding Map Projections: GIS by ESRI. ESRI Press. https://gis.icao.int/icaoetod/map_projections[1].pdf

Kerr, J. T., & Deguise, I. (2004). Habitat loss and the limits to endangered species recovery. Ecology Letters, 7(12), 1163–1169. https://doi.org/10.1111/j.1461-0248.2004.00676.x

Komonen, A. (2007). Are we conserving peripheral populations? An analysis of range structure of longhorn beetles (Coleoptera: Cerambycidae) in Finland. Journal of Insect Conservation, 11(3), 281–285. https://doi.org/10.1007/s10841-006-9043-8

Kraus, D., & Hebb, A. (2020). Southern Canada’s crisis ecoregions: identifying the most significant and threatened places for biodiversity conservation. Biodiversity and Conservation, 29(13), 3573–3590. https://doi.org/10.1007/s10531-020-02038-x

Kraus, D., Murphy, S., & Armitage, D. (2021). Ten bridges on the road to recovering Canada’s endangered species. Facets, 6, 1088–1127. https://doi.org/10.1139/facets-2020-0084

Lenth, R. V. (2016). Least-squares means: The R package lsmeans. Journal of Statistical Software, 69(1). https://doi.org/10.18637/jss.v069.i01

Littlefield, C. E., Krosby, M., Michalak, J. L., & Lawler, J. J. (2019). Connectivity for species on the move: supporting climate-driven range shifts. Frontiers in Ecology and the Environment, 17(5), 270–278. https://doi.org/10.1002/fee.2043

Maxwell, S. L., Cazalis, V., Dudley, N., Hoffmann, M., Rodrigues, A. S. L., Stolton, S., Visconti, P., Woodley, S., Kingston, N., Lewis, E., Maron, M., Strassburg, B. B. N., Wenger, A., Jonas, H. D., Venter, O., & Watson, J. E. M. (2020). Area-based conservation in the twenty-first century. Nature, 586(7828), 217–227. https://doi.org/10.1038/s41586-020-2773-z

Mooers, A. O., Doak, D. F., Scott Findlay, C., Green, D. M., Grouios, C., Manne, L. L., Rashvand, A., Rudd, M. A., & Whitton, J. (2010). Science, policy, and species at risk in Canada. BioScience, 60(10), 843–849. https://doi.org/10.1525/bio.2010.60.10.11

Natural Earth. (2020). Natural earth vector countries. Version 4.1.0 [online]: Available from https://naturalearthdata.com

Orme, C. D. L., Davies, R. G., Burgess, M., Eigenbrod, F., Pickup, N., Olson, V. A., Webster, A. J., Ding, T. S., Rasmussen, P. C., Ridgely, R. S., Stattersfield, A. J., Bennett, P. M., Blackburn, T. M., Gaston, K. J., & Owens, I. P. F. (2005). Global hotspots of species richness are not congruent with endemism or threat. Nature, 436(7053), 1016–1019. https://doi.org/10.1038/nature03850

Parks Canada. (2019). Parks Canada attendance 2019-20. Parks Canada attendance. https://publications.gc.ca/site/eng/9.839688/publication.html

Prendergast, J. R., Quinn, R. M., Lawton, J. H., Eversham, B. C., & Gibbons, D. W. (1993). Rare species, the coincidence of diversity hotspots and conservation strategies. Nature, 365(6444), 335–337. https://doi.org/10.1038/365335a0

QGIS Development Team. (2021). QGIS Geographic Information System. Available from http://qgis.osgeo.org.

R Core Team. (2021). R: A language and environment for statistical computing. R Foundation for Statistical Computing, Vienna, Austria. Available from https://www.R-project.org/.

Raymond, C. V., Wen, L., Cooke, S. J., & Bennett, J. R. (2018). National attention to endangered wildlife is not affected by global endangerment: a case study of Canada’s species at risk program. Environmental Science and Policy, 84(August 2017), 74–79. https://doi.org/10.1016/j.envsci.2018.03.001

Reid, W. V. (1998). Biodiversity hotspots. Trends in Ecology and Evolution, 13(7), 275–280. https://doi.org/10.1016/S0169-5347(98)01363-9

Rodrigues, A. S. L., & Gaston, K. J. (2002). Rarity and conservation planning across geopolitical units. Conservation Biology, 16(3), 674–682. https://doi.org/10.1046/j.1523-1739.2002.00455.x

Rotenberry, J. T., & Balasubramaniam, P. (2020). Connecting species’ geographical distributions to environmental variables: range maps versus observed points of occurrence. Ecography, 43(6), 897–913. https://doi.org/10.1111/ecog.04871

Thomas, C. D., Gillingham, P. K., Bradbury, R. B., Roy, D. B., Anderson, B. J., Baxter, J. M., Bourne, N. A. D., Crick, H. Q. P., Findon, R. A., Fox, R., Hodgson, J. A., Holt, A. R., Morecroft, M. D., O’Hanlon, N. J., Oliver, T. H., Pearce-Higgins, J. W., Procter, D. A., Thomas, J. A., Walker, K. J., … Hill, J. K. (2012). Protected areas facilitate species’ range expansions. Proceedings of the National Academy of Sciences of the United States of America, 109(35), 14063–14068. https://doi.org/10.1073/pnas.1210251109

Titley, M. A., Snaddon, J. L., & Turner, E. C. (2017). Scientific research on animal biodiversity is systematically biased towards vertebrates and temperate regions. PLoS ONE, 12(12), 1– https://doi.org/10.1371/journal.pone.0189577

Vellend, M. (2003). Habitat loss inhibits recovery of plant diversity as forests regrow. Ecology, 84(5), 1158–1164. https://doi.org/10.1890/0012-9658(2003)084[1158:HLIROP]2.0.CO;2

Venter, O., Fuller, R. A., Segan, D. B., Carwardine, J., Brooks, T., Butchart, S. H. M., Di Marco, M., Iwamura, T., Joseph, L., O’Grady, D., Possingham, H. P., Rondinini, C., Smith, R. J., Venter, M., & Watson, J. E. M. (2014). Targeting global protected area expansion for imperiled biodiversity. PLoS Biology, 12(6). https://doi.org/10.1371/journal.pbio.1001891

Venter, O., Magrach, A., Outram, N., Klein, C. J., Possingham, H. P., Marco, M. Di, & Watson, J. E. M. (2017). Bias in protected-area location and its effects on long-term aspirations of biodiversity conventions. Conservation Biology, 32(1), 127–134. https://doi.org/10.1111/cobi.12970

Watson, J. E. M., Dudley, N., Segan, D. B., & Hockings, M. (2014). The performance and potential of protected areas. Nature, 515(7525), 67–73. https://doi.org/10.1038/nature13947

Wells, J. V., Robertson, B., Rosenberg, K. V., & Mehlman, D. W. (2010). Global versus local conservation focus of U.S. state agency endangered bird species lists. PLoS ONE, 5(1), 3–7. https://doi.org/10.1371/journal.pone.0008608

White, M. P., Alcock, I., Grellier, J., Wheeler, B. W., Hartig, T., Warber, S. L., Bone, A., Depledge, M. H., & Fleming, L. E. (2019). Spending at least 120 minutes a week in nature is associated with good health and wellbeing. Scientific Reports, 9(1), 1–11. https://doi.org/10.1038/s41598-019-44097-3

Woo-Durand, C., Matte, J. M., Cuddihy, G., McGourdji, C. L., Venter, O., & Grant, J. W. A. (2020). Increasing importance of climate change and other threats to at-risk species in Canada. Environmental Reviews, 28(4), 449–456. https://doi.org/10.1139/er-2020-0032

Yong, D. L., Barton, P. S., Okada, S., Crane, M., & Lindenmayer, D. B. (2016). Birds as surrogates for mammals and reptiles: are patterns of cross-taxonomic associations stable over time in a human-modified landscape? Ecological Indicators, 69, 152–164. https://doi.org/10.1016/j.ecolind.2016.04.013

